# Effects of lactate, super-GDF9 and low oxygen tension during biphasic in vitro maturation on the bioenergetic profiles of mouse cumulus-oocyte-complex

**DOI:** 10.1101/2022.11.09.514870

**Authors:** Nazli Akin, Gamze Ates, Lucia von Mengden, Anamaria-Cristina Herta, Cecilia Meriggioli, Katy Billooye, William A. Stocker, Brecht Ghesquiere, Craig A. Harrison, Wilfried Cools, Fabio Klamt, Ann Massie, Johan Smitz, Ellen Anckaert

**Author notes:** CORRESPONDING AUTHOR: Correspondences should be addressed to Nazli Akin – – Follicle Biology Laboratory, VUB Jette Campus, Laarbeeklaan 103, 1090, Brussels. **AUTHORS’ ROLES** NA: Designed the experiments, analyzed the data, prepared the manuscript. NA, AH, KB: Performed cultures and in vitro fertilization experiments, collected samples for enzymatic assays. NA, GA: Performed mitochondrial function tests with Seahorse Analyzer and interpreted the data. NA, AH, CM: Performed enzymatic assays. NA, BG: Performed stainings, imaging and image analysis. LVM: Provided supervision on the enzymatic assays. WS, CH: Synthesized the super-GDF9. WC: Provided support on statistics and performed statistical analysis. FK, AM: Provided supervision on the data interpretation. JS, EA: Supervised the project and secured the funding. All authors have read the manuscript and agreed on the final version. **CONFLICT OF INTEREST** Authors report no conflict of interest. **FUNDING:** This project was funded by the Fonds voor Wetenschappelijk Onderzoek Vlaanderen (FWO) Excellence of Science (EOS, FWO-F.R.S-FNRS; G0F3118N), FWO medium-scale research infrastructure program (I001420N) and the Vrije Universiteit Brussel (OZR). GA is supported by FWO (12B3223N).

## Abstract

In vitro maturation (IVM) is an alternative assisted reproductive technology (ART) with reduced hormone related side-effects and treatment burden compared to conventional IVF. Capacitation (CAPA)-IVM is a biphasic IVM system with improved clinical outcomes compared to standard monophasic IVM. Yet, CAPA-IVM efficiency compared to conventional IVF is still suboptimal in terms of producing utilizable blastocysts. Previously we have shown that CAPA-IVM leads to a precocious increase in cumulus cell (CC) glycolytic activity during cytoplasmic maturation. In the current study, considering the fundamental importance of CCs for oocyte maturation and cumulus-oocyte complex (COC) microenvironment, we further analyzed the bioenergetic profiles of maturing CAPA-IVM COCs. Through a multi-step approach, we (i) explored mitochondrial function of the in vivo and CAPA-IVM matured COCs through real-time metabolic analysis with Seahorse analyzer; and to improve COC metabolism (ii) supplemented the culture media with lactate and/or super-GDF9 (an engineered form of growth differentiation factor 9) and (iii) reduced culture oxygen tension. Our results indicated that the pre-IVM step is delicate and prone to culture related disruptions. Lactate and/or super-GDF9 supplementations failed to eliminate pre-IVM induced stress on COC glucose metabolism and mitochondrial respiration. However, when performing pre-IVM culture under 5% oxygen tension, CAPA-IVM COCs showed a similar bioenergetic profiles compared to in vivo matured counterparts. This is the first study providing real-time metabolic analysis of the COCs from a biphasic IVM system. The currently used analytical approach provides the quantitative measures and the rational basis to further improve IVM culture requirements.

## INTRODUCTION

In vitro fertilization (IVF) involves the stimulation of ovaries with exogenous gonadotropins for several days, followed by the triggering of ovulation (e.g. with human chorionic gonadotropin (hCG) administration). In vitro maturation (IVM), on the other hand, is an alternative assisted reproductive technology (ART), mostly offered to patients with polycystic ovarian syndrome (PCOS), where immature cumulus-oocyte-complexes (COCs) are collected from small/mid-antral follicles and matured in vitro. Patient preparation for IVM requires minimal (2-3 days) or no gonadotropin stimulation, and no hCG priming is necessary [1,2]. Therefore, it is a mild and patient friendly approach with reduced risks of hormone-related side effects and lower treatment costs per cycle.

Earlier research has shown that standard IVM systems are leading to spontaneous oocyte nuclear maturation, upon removal of the COC from the inhibitory follicular environment by a lack of intrinsic factors regulated by somatic cell compartment and loss of cumulus-oocyte connections [3]. This precipitous resumption of meiosis is however not accompanied with cytoplasmic maturation (or capacitation). In order to remediate, bi-phasic IVM systems, with a pre-IVM step, allowing oocyte capacitation prior to triggering nuclear maturation have been developed [4–8]. CAPA-IVM, being one successful example of a bi-phasic IVM system, uses C-type natriuretic peptide (CNP) during the pre-IVM step to regulate cAMP concentrations to block oocyte meiosis resumption, followed by an IVM step with epidermal growth factor network peptides and FSH to induce nuclear maturation [9,10]. Bi-phasic CAPA-IVM has improved the competence of the oocytes derived from small antral follicles and led to increased oocyte maturation and clinical pregnancy rates [10–14] through synchronized nuclear and cytoplasmic maturation [9,12]. Even though CAPA-IVM is superior to the standard IVM systems, a current challenge to reduce the high attrition rate from mature oocytes to utilizable blastocysts, which are still lower in number compared to conventional IVF [14]. One potential solution for reducing the efficiency gap between CAPA-IVM and conventional IVF could be through optimizing the culture conditions (such as media formulation) to mimic the in vivo environment, thus mitigating the negative effects inflicted on the oocytes by in vitro culture [3].

The follicular microenvironment depends on several energy substrates including glucose, lactate, and pyruvate, in varying concentrations as well as oxygen [15]. For obtaining a competent oocyte, proper utilization of follicular fluid (FF) nutrients and oxygen are important as much as the substrate availability [16]. During in vivo maturation, cumulus cells (CCs) take up the glucose from FF and perform glycolysis to provide the oocytes with pyruvate [17–19], which is then metabolized by oocytes through mitochondrial oxidative phosphorylation (OXPHOS) for ATP generation [17,20,21]. FF also supplies sufficient levels of oxygen to the COCs residing in an avascular environment [22,23] for energy transformation through pyruvate oxidation [24,25]. In fact, it was shown that the oocyte is the main receiver of the FF oxygen, as only up to 5% of this essential gas is absorbed by CCs within the COCs [23,26–28]. Like glucose, lactate in the medium is metabolized by the CCs to produce pyruvate which can support the oocyte during maturation [29]. Furthermore, mouse oocytes can metabolize culture media pyruvate, but not lactate-derived pyruvate, to convert into energy in mitochondria [30]. Nevertheless, oxidation of pyruvate is crucial for oocytes regardless of the source it has been derived from [31].

Alterations in metabolic activity are known culture-induced negative effects, and IVM-inflicted perturbations in COC metabolism have been observed in several species [32–36]. Recently, we have shown an impaired precocious increase in glycolytic activity in CCs during the pre-IVM phase of the mouse CAPA-IVM when compared to their in vivo matured counterparts [37], which may contribute to the lower in vitro oocyte competency. Consequently, limiting pre-IVM glycolysis and/or focusing on improving CC function might possibly enhance CAPA-IVM culture outcomes. The former could be potentially achieved through supplementing lactate to CAPA-IVM basal media, which was missing in the culture medium used in the former study. Lactate is the end-product of lactic acid fermentation, with a known role in inhibiting the activity of the glycolytic enzyme phosphofructokinase (PFK) in several tissues [38]. In early-stage murine embryos, lactate also reduces glycolytic activity indirectly through its conversion into pyruvate [39]. On the other hand, CC function could be improved through supplementing IVM media with oocyte-secreted factors (OSFs). OSFs enhance oocyte quality through orchestrating several CC metabolic functions; including glycolysis, amino acid uptake and cholesterol biosynthesis (reviewed in [40,41]). CAPA-IVM media supplementation with cumulin, a heterodimer of bone morphogenic protein-15 (BMP15) and growth differentiation factor-9 (GDF9), regulates the expression levels of ovulatory cascade genes to a level more resembling that of in vivo matured CCs [42]. Similarly, the engineered OSF super-GDF9, which has over 1000-fold greater bioactivity than wild-type GDF9 [43], visibly improved COC mucification and extracellular matrix (ECM) elasticity in CAPA-IVM which was more comparable to the ECM morphology of in vivo matured COCs [42].

Regulating oxygen tension of the culture environment could be another possible approach to improve COC metabolism. While there is a long-standing consensus on the requirement of low oxygen tension during in vitro embryo culture, the optimal oxygen concentration for IVM is still controversial. The low oxygen (2%-9%) in the in vivo COC environment [44,45] is overridden in the in vitro settings where oxygen concentrations are at atmospheric levels. Oocyte mitochondria use substrates produced by CC glycolysis to perform OXPHOS that will provide the necessary energy throughout oocyte maturation, fertilization, and preimplantation development (reviewed in [41]). Supraphysiological oxygen levels could trigger mitochondrial activity and lead to imbalanced ATP generation [46], as well as the accumulation of reactive oxygen species (ROS). Moreover, immature mitochondria in GV oocytes might not be equipped to withstand the high oxygen environment during pre-IVM. Thus, regulating the oxygen tension during IVM to levels comparable to those in vivo could also offer a way of by-passing the CC metabolism and targeting the oocyte OXPHOS directly, and be beneficial for sustaining COCs innate bioenergetic profile.

While there was no indication for a faulty oocyte metabolism in our previous spectrophotometric analysis-based study, it was also lacking direct insights on OXPHOS and mitochondrial respiration [37]. Hence in this study, we initially studied the mitochondrial function of in vivo and CAPA-IVM matured COCs, in order to gain a better understanding of the possible adverse effects of defective CC glycolysis on the ATP production of the oocytes via OXPHOS. Then, we hypothesized that supplementing CAPA-IVM media with lactate and super-GDF9, either separately or in combination, could restore the defective CAPA-IVM CC glucose metabolism and basal respiration through limiting glycolysis and/or improving CC function. Finally, we analyzed the effect of low oxygen tension during pre-IVM culture on oocyte competence and mitochondrial function. This is the first study applying an extracellular flux analysis to understand the changes in the respiration profile of COCs during a bi-phasic IVM system. Our results suggest that low oxygen (5%) culture environment during cytoplasmic maturation might possibly be the key of improving CAPA-IVM success.

## MATERIALS AND METHODS

### Animal model

Animals used in this study were the F1 mice of C57BL/6j x CBA/ca hybrid (Charles River Laboratories, France). All mice were housed and bred with the approval of Vrije Universiteit Brussel’s local ethical committee (approval no: 21-216-1) and in accordance with the Belgian Legislation for animal care.

### COC collection

Ovaries were collected from 19- to 21-day-old prepubertal female mice in Leibovitz L15 (Sigma-Aldrich, Belgium) supplemented with 10% fetal bovine serum (FBS), 100 IU/ml penicillin and 100 μg/ml streptomycin (all from Thermo Fisher Scientific, Belgium). This study included three types of in vivo samples: (i) baseline-COCs (baseline), retrieved in mice that received no hormonal stimulation; (ii) in vivo GV-COCs, collected 48 h after stimulation with 2.5 IU Folligon (Intervet, Netherlands); and (iii) in vivo MII-COCs, collected after the stimulation with 2.5 IU Folligon for 48 h followed by 2.5 IU Chorulon (Intervet) for 14 h. Mice from the second and the third groups are considered as in vivo super-ovulated (SO) mice. Different from in vivo samples, CAPA-IVM COCs were collected from unstimulated mice that did not receive any hormonal stimulation. Collection medium was supplemented with 0.2 mM IBMX (3-isobutyl-1-methylxanthine; Sigma-Aldrich) to prevent meiotic resumption prior to culture. COCs for CAPA-IVM culture and in vivo baseline and GV-COCs were collected from the harvested ovaries, through puncturing small antral follicles with insulin needles. In vivo MII-COCs were collected from the fallopian tubes.

### CAPA-IVM culture

Basal culture medium for CAPA-IVM was prepared as described previously [9]. Briefly, alpha-MEM (Thermo Fisher Scientific) was supplemented with 2.5 % FBS, 5 ng/ml sodium selenite, 5 μg/mL apo-transferrin, 5 ng/ml insulin, 10 nM 17-beta estradiol (E2) (all from Sigma-Aldrich) and 2.5 mIU/ml FSH (Merck, Belgium). Basal media was supplemented with 25 nM CNP-22 (Phoenix Europe, Germany) for pre-IVM culture and 50 ng/mL recombinant mouse epiregulin (EREG; Bio-techne, United Kingdom) for IVM culture. Individual COCs were cultured for 48 h in pre-IVM media, followed by 18 h in IVM media, in 96-well round bottom ultra-low attachment plates (Corning) at 37 ^O^C, 5% CO_2_ and 100% humidity.

### Evaluation of MII oocyte rate and oocyte competence

At the end of the IVM culture, COCs were processed for evaluation of several different end points. For one set of experiments, COCs were denuded mechanically to assess the maturation rate and measure MII oocyte diameters. Maturation rate was calculated by dividing the number of oocytes with a visible polar body to the total number of denuded oocytes. For another set of experiments, oocyte competence was assessed through performing IVF and embryo culture (details were explained in [42]). In IVF experiments, cleavage rate on day 2 (D2) and blastocysts rates on D5 were assessed as end points. Cleavage rate was calculated over the total number of inseminated COCs, whereas D5 blastocyst rate was calculated over the total number of cleaved embryos on D2.

### Enzymatic assays: Sample preparation and assays

For the enzymatic assays, in vitro matured GV and MII COCs were collected at the end of pre-IVM and IVM cultures, respectively. Mechanically denuded oocytes and their corresponding cumulus cells were collected in pools of five (one biological replicate), snap frozen and stored at −80 ^O^C. Cell lysis was performed as described before [37] using an in-house prepared lysis buffer and through applying three freeze-thaw cycles. Protein content was measured using a modified Bradford assay [47]. Samples were aliquoted and stored at −80 ^O^C until performing the assays. Phosphofructokinase (PFK) (E.C.: 2.7.1.11) and lactate dehydrogenase (LDH) (E.C.: 1.1.1.27) activities and concentrations of pyruvate and lactate were measured in samples via enzymatic assays (all from Sigma-Aldrich). All assays have been optimized for the current sample types as explained in the previous studies [37,48] and specific protocol settings for each assay can be found in [37]. All assays were performed on a SpectraMax i3 (Molecular Devices) using appropriate 96-well plates.

### Extracellular Flux Analysis with Seahorse Analyzer

The mitochondrial function of in vitro (with CAPA-IVM) and in vivo (baseline and SO) grown and matured COCs were studied using the Seahorse metabolic flux analyzer (XFe96) and the Mito Stress Test (Agilent Technologies). The decision to analyze COCs as a whole complex rather than focusing on the individual cell types was made based on the proven essential bi-directional communication between CCs and oocytes [33,41]. Nevertheless, as 95% of the oxygen is directed to cumulus-enclosed-oocytes [49], real-time OCR of COCs can be used as a straightforward indicator of functionality of oocyte mitochondria. The Mito Stress Test allows for the study of mitochondrial function in real-time, through serial injection of several inhibitors of the electron transport chain and recording the oxygen consumption rate (OCR) of the cells. Basal cellular respiration, as well as mitochondrial respiration, maximal respiration, proton leak, ATP-coupled respiration, non-mitochondrial oxygen consumption, spare respiratory capacity and coupling efficiency are calculated using Seahorse Analytics (seahorseanalytics.agilent.com). Definitions of the reported endpoints are provided in Table 1.

**Table 1:**
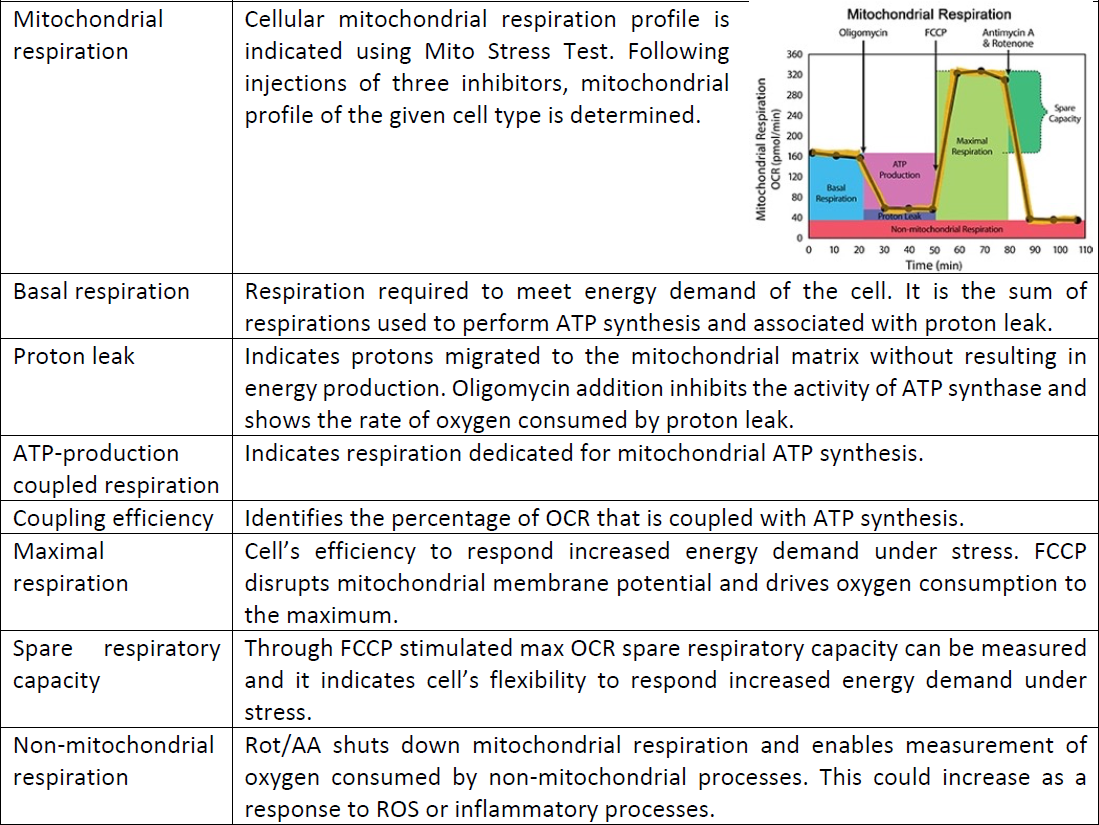
Explanation of Mito Stress Test assay parameters [49].

|  |  |
| --- | --- |
| Mitochondrial respiration | Cellular mitochondrial respiration profile is indicated using Mito Stress Test. Following injections of three inhibitors, mitochondrial profile of the given cell type is determined. |
| Basal respiration | Respiration required to meet energy demand of the cell. It is the sum of respirations used to perform ATP synthesis and associated with proton leak. |
| Proton leak | Indicates protons migrated to the mitochondrial matrix without resulting in energy production. Oligomycin addition inhibits the activity of ATP synthase and shows the rate of oxygen consumed by proton leak. |
| ATP-production coupled respiration | Indicates respiration dedicated for mitochondrial ATP synthesis. |
| Coupling efficiency | Identifies the percentage of OCR that is coupled with ATP synthesis. |
| Maximal respiration | Cell's efficiency to respond increased energy demand under stress. FCCP disrupts mitochondrial membrane potential and drives oxygen consumption to the maximum. |
| Spare respiratory capacity | Through FCCP stimulated max OCR spare respiratory capacity can be measured and it indicates cell's flexibility to respond increased energy demand under stress. |
| Non-mitochondrial respiration | Rot/AA shuts down mitochondrial respiration and enables measurement of oxygen consumed by non-mitochondrial processes. This could increase as a response to ROS or inflammatory processes. |

The XFe96 sensor cartridge was hydrated overnight with sterile water in a CO_2_ free incubator at 37°C. Following the first hydration step, water was replaced with calibrant solution (preincubated overnight at 37°C in a CO_2_-free incubator) and the plate placed back into the CO_2_ free incubator setting (at 37°C) for 1 hour. Meanwhile, Seahorse plates were coated with Cell-Tak (Corning) following the basic coating protocol of the manufacturer. COCs were collected at the end of pre-IVM and IVM and washed three times in droplets of assay medium (prepared by supplementing XF DMEM with 1 mM pyruvate, 2 mM glutamine and 5 mM glucose [50]) before transfer into the Seahorse plate. Similarly, baseline, in vivo GV- and MII-COCs were collected following the same preparation protocol as described above. 5-15 COCs (one biological replicate) were transferred per assay well together with 20 μl XF assay medium (final assay volume per well is 180 μl), and the plate was placed into the CO_2_ free incubator at 37°C to de-gas for 1 hour. Final concentrations (per well) for Mito Stress Test Kit reagents’ oligomycin, FCCP (carbonyl cyanide p-trifluoro-methoxyphenyl hydrazone) and Rot/AA (rotenone/antimycin A) were 1 μM, 2.5 μM and 2.5 μM, respectively [50]. Upon completing the runs with the Seahorse, the contents of each well were lysed using 75 μl extraction buffer (2% sodium dodecyl sulfate, 60 nM Tris, 100 mM DTT, protease and phosphatase inhibitor cocktails) [51]. Supernatants were collected after incubating the samples at 37°C for 30 mins followed by centrifuging at 4°C at 10,000 rpm for 15 mins. Protein concentration per sample (μg/μl) was calculated using Qubit Protein Assay (Thermo Fisher) and sample protein concentration was used to normalize the data.

Extracellular flux analysis experiments were completed performing 4 repeats (only three were including samples from CAPA-IVM low O_2_ culture). In total 82 female mice were used to collect COCs for CAPA-IVM culture, 11 female mice were used to collect COCs at baseline and 16 female mice were stimulated to collect SO GVs-(8 mice) or MII-(8 mice) COCs.

### CellROX and JC-1 staining, imaging and image analysis

CellROX Deep Red (Thermo Fisher) was used to measure the intracellular ROS induced oxidative stress. Mitochondrial membrane potential was calculated using the MitoProbe JC-1 assay kit (Thermo Fisher). All media and oil were pre-equilibrated at 37°C, 5% CO_2_ and 100% humidity prior to use. Immunofluorescence labeling was carried out in 25 μl drops of staining solution (5 μM CellRox and 2 μM JC-1 in M16 medium) under oil. GV oocytes were mechanically denuded, transferred to the staining dish (one condition per droplet) and incubated for 30 mins at 37°C, 5% CO_2_ and 100% humidity. Following three washing steps in M16 medium, oocytes were imaged in glass bottom confocal dishes (MatTek) in 2 μl dops of M16, under oil, on a Zeiss LSM 800 confocal microscope using live imaging chamber (37°C, 5% CO_2_, and humidity).

Three images with a z-step of 10 μm were taken on a 40x magnification. Accumulated ROS signal was detected at 653 nm (red). JC-1 aggregates, indicating active mitochondria membrane potential was detected at 565 nm (orange), while JC-1 monomers reflective of electrochemical potential loss was measured at 493 nm (green). Fluorescence signal was measured for each channel, on all z-stacks, within the region of interest (in the ooplasm) as the mean intensity value using the image analysis wizard in ZenBlue (Zeiss software). The intensity values from three stacks were averaged as an individual value for each sample. Averaged intensity values were normalized against average intensity obtained in CAPA-IVM control. Overall mitochondrial membrane potential per GV oocyte was calculated as the aggregate to monomer ratio (orange/green) detected in the ooplasm [52].

Experiments for stainings were completed performing 3 repeats, collecting COCs from 4 stimulated and 52 unstimulated mice (baseline and cultured). Total numbers of imaged GV oocytes are as follows: Baseline: 42; SO-GV: 44, CAPA-IVM Control (20% O_2_): 34, CAPA-IVM Lactate only: 27, CAPA-IVM GDF9 only: 24, CAPA-IVM Lactate+GDF9: 26, CAPA-IVM Control (5% O_2_): 26.

### Experimental Design

This study consisted of three parts, as illustrated in Figure 1. Designs of each experiment, as well as assessed end points, are explained below.

**Figure 1:**
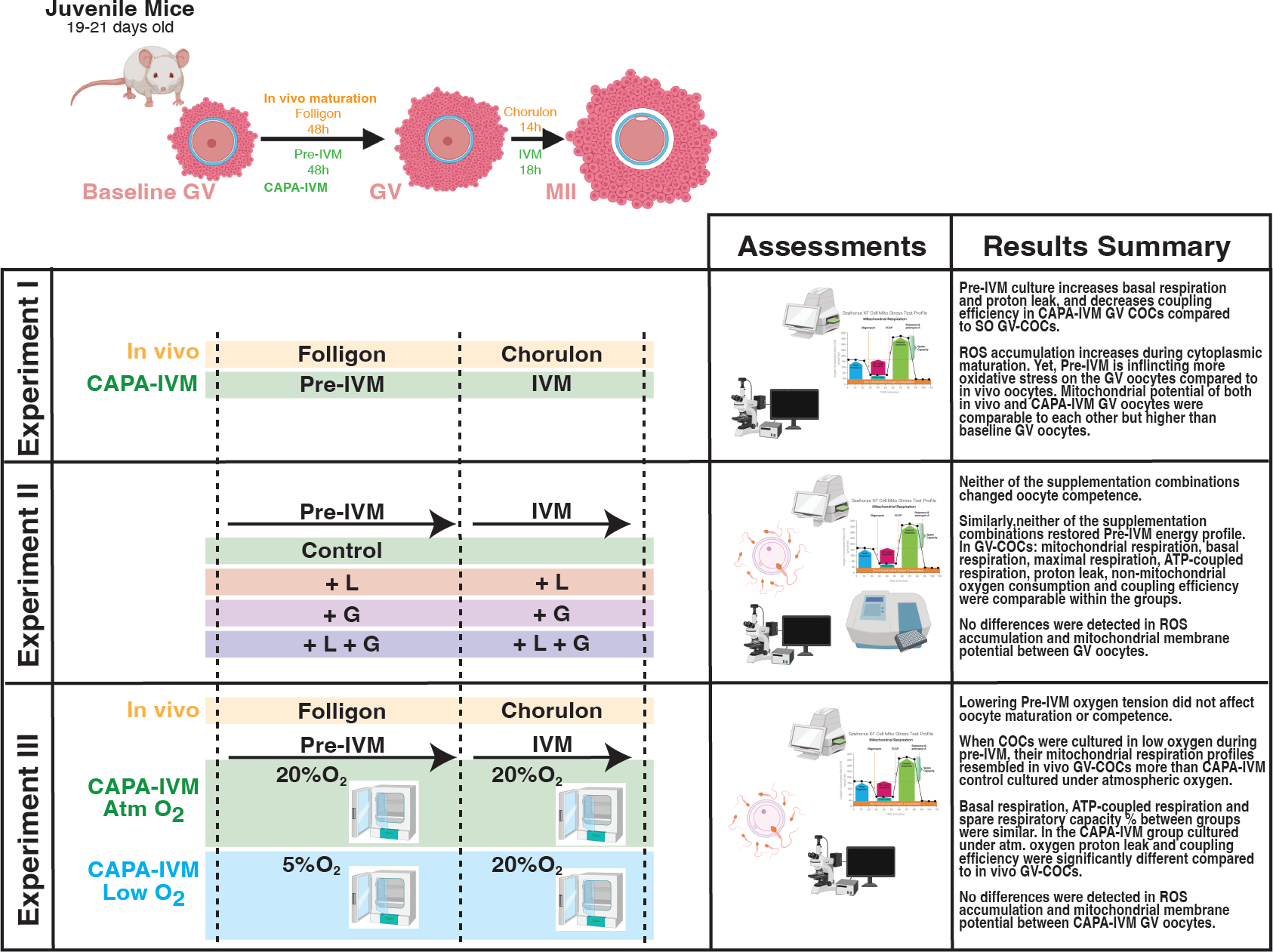
Illustration of experimental design and brief summary of results. In Experiment I, mitochondrial function (Seahorse Analyzer, confocal imaging) of in vivo and CAPA-IVM matured COCs was studied. Experiment II focused on individual (enzymatic assays, IVF, Seahorse Analyzer) and combined effects (IVF, Seahorse Analyzer, confocal imaging) of lactate and super-GDF9 supplementation to CAPA-IVM culture media. In Experiment III, effect of low oxygen tension during pre-IVM on oocyte competence (IVF) and COC metabolism (Seahorse Analyzer, confocal imaging) was assessed. L: Lactate. G: Super-GDF9. Atm: Atmospheric. Figure was created using BioRender.

#### Experiment I: Mitochondrial function in the in vivo matured vs CAPA-IVM COCs

As a follow-up to the previously published results [37], mitochondrial function of the in vivo and in vitro matured COCs was assessed using the Seahorse metabolic flux analyzer and Cell-ROX staining. Furthermore, mitochondrial membrane potential was assessed through JC-1 staining. In vivo matured COCs and CAPA-IVM COCs were obtained as described above following hormonal stimulations and after standard CAPA-IVM culture. In order to track changes taking place during oocyte capacitation (following Folligon injection in vivo, during pre-IVM in vitro) a baseline in vivo condition was also included.

#### Experiment II: Individual and combined effects of lactate and super-GDF9 supplementation on CAPA-IVM COC metabolism

The individual effect of lactate on CAPA-IVM COC metabolism was studied trough supplementing the basal culture media with Calcium L-lactate hydrate (Sigma-Aldrich). An initial dose finding study was performed, during which the effect of two doses of lactate (1 mM and 2 mM) - added at the pre-IVM step-was evaluated on the MII rate and oocyte size. Dose selection for this initial dose finding study was based on literature[53]. No disruption in the system was detected with either of the doses (Supplementary Figure 1A and B). Given the acidity of the environment affects oocyte maturation, and lactate concentration decreases in the reproductive tract during maturation [15], the effect of lactate in our CAPA-IVM system was tested through addition either to pre-IVM only (2 mM) or to both culture steps where lactate concentration was decreased during IVM (pre-IVM 2 mM and IVM 1 mM). End points were assessment of the MII oocyte rate, oocyte competence and performing enzymatic assays.

The individual effect of super-GDF9 on CAPA-IVM COC metabolism was studied by supplementing basal culture media with super-GDF9 (50 ng/ml in pre-IVM and 25 ng/ml in IVM). We have previously demonstrated that given doses of OSF supplementation of our CAPA-IVM system during both culture steps led to enhanced mucification and elasticity of the extracellular matrix in COCs, a feature that was in-line with cumulus cell gene expression results [42]. Thus, we applied the same culture protocol in the current study. We used super-GDF9 in the current study given its higher potency and purer chemical composition compared to cumulin [43]. Preparation of the engineered super-GDF9 is explained in detail in [43] and [42]. Given that oocyte competence and MII rates following super-GDF9 supplementation have already been assessed and reported in a previous study and concluded to be comparable [42], cultures from this condition were utilized to perform enzymatic assays.

Following the identification of the individual effects of lactate and super-GDF9, their combined effect was tested against individual supplementations. For combined testing, the pre-IVM culture step was supplemented with 2 mM lactate and 50 ng/ml super-GDF9 followed by the IVM culture step with 1 mM lactate and 25 ng/ml super-GDF9. Cultures from this experimental condition were performed to assess MII oocyte rate, MII oocyte size, oocyte competence, mitochondrial function, ROS accumulation and mitochondrial membrane potential.

#### Experiment III: Effect of oxygen tension

As a follow-up set of experiments, the effect of culture environment oxygen concentration was assessed. Considering the collective results obtained from experiments I and II, we focused on adjusting the oxygen tension during the pre-IVM step. For the control group, standard CAPA-IVM culture settings, as described above, were followed. For the experimental group (low oxygen CAPA-IVM), the COCs were cultured at 37 ^O^C, 5% CO_2_, 6% O_2_ and 100% humidity during pre-IVM and 37 ^O^C, 5% CO_2_, atmospheric O_2_ and 100% humidity during IVM. SO COCs were also included for assessment of mitochondrial function with the Seahorse Mito Stress Test. MII oocyte rate and oocyte competence, as well as mitochondrial function, ROS levels and mitochondrial membrane potential were assessed and compared between groups.

### Statistical analysis

Maturation, D2 cleavage and blastocyst rates were arcsine transformed prior to statistical analysis with one-way ANOVA followed by Tukey’s post hoc test. For the enzymatic assays, all relevant groups were compared through ordinary one-way ANOVA using Sidak‘s correction on log_2_ transformed data. For the Mito Stress Test, the data was analyzed using mixed linear models to accommodate the correlations following the clustered data, making use of each observation (instead of averages) collected from the independent experiments. For CellROX and JC-1 staining, values collected from each experimental condition were normalized to the average value of the CAPA-IVM control GV oocytes, and the statistical difference was calculated either with one-way ANOVA (Experiments 1&2; unpaired; Sidak’s correction) or with paired t-test (Experiment III). For all tests, significance was considered when p<0.05 and confidence interval was 95%. Statistical analyses were performed using GraphPad Prism (GraphPad Software, La Jolla, California, USA, www.graphpad.com) and R (R Foundation for Statistical Computing, Vienna, Austria).

## RESULTS

### Respiration patterns of immature SO and CAPA-IVM COCs are dramatically different

Mitochondrial respiration data highlight that cytoplasmic (Figure2A) and nuclear (Figure2B) maturation follow distinct patterns, as well as in vivo and in vitro maturation. Compared to baseline, both basal and maximal respiration decreases significantly in SO GV-COCs following folligon injection (Figure2C, p<0.05; Figure2D, p<0.001). On the other hand, basal and maximal respiration of CAPA-IVM GV-COCs is comparable to the baseline COCs, showing a different pattern compared to their SO counterparts. This difference between in vivo and in vitro GV-COCs is also evident, as basal and maximal respiration rates of the two groups are significantly different (Figure2C,p<0.001; Figure2D, p<0.01). While there is an increasing pattern (although not significant) in both maximal and basal respiration during in vivo maturation (GV to MII transition), in CAPA-IVM both decrease with significant differences observed for COCs after IVM (Figure2C, p<0.05; Figure2D, p<0.01).

**Figure 2:**
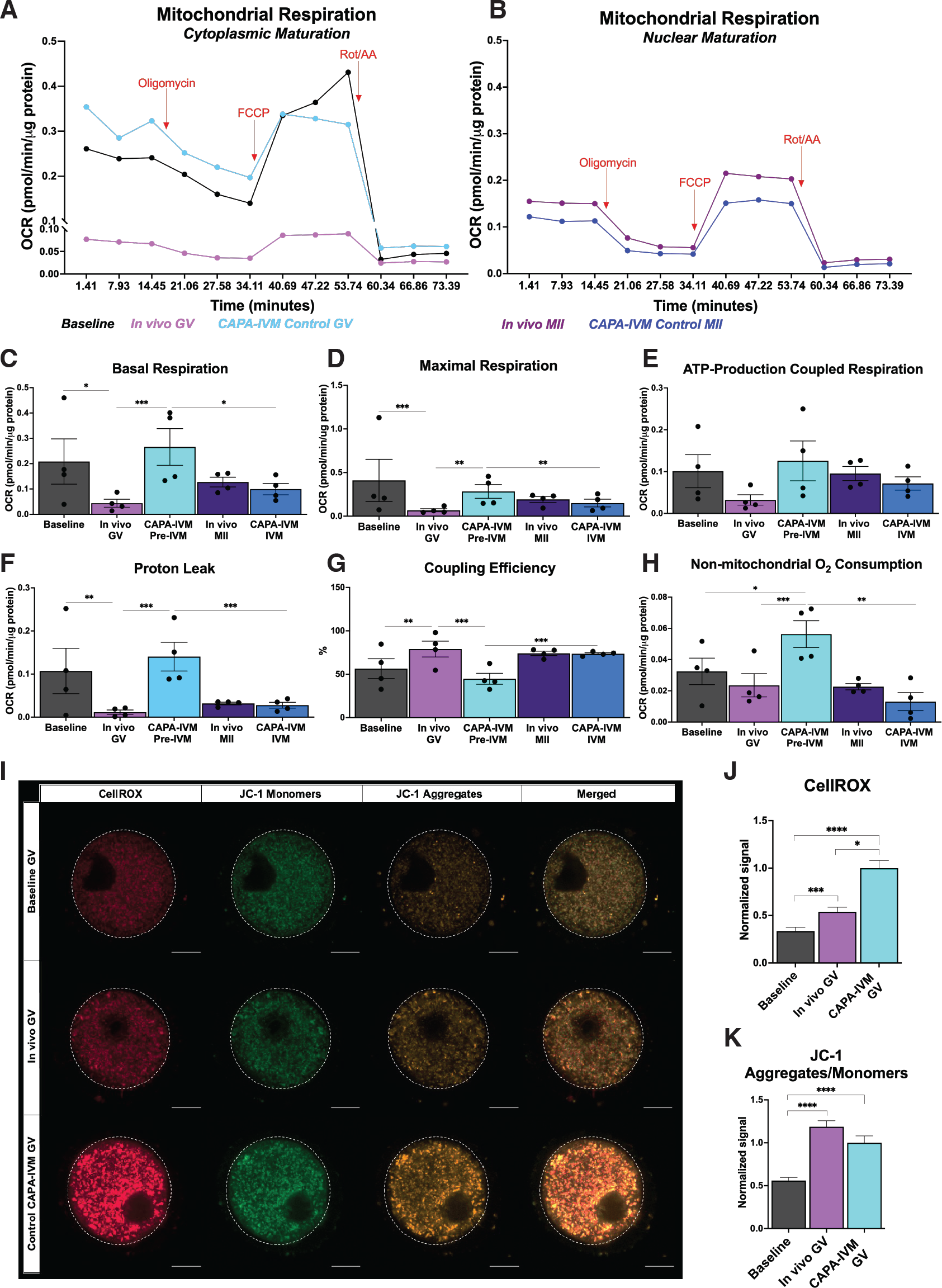
Mitochondrial function, ROS accumulation and mitochondrial membrane potential were compared between in vivo vs CAPA-IVM matured COCs. **(A-H)** show results obtained through real-time metabolic analysis with Mito Stress Test (Seahorse Analyzer). **(I-K)** Panels illustrates CellROX (red), JC-1 monomer (green) and JC-1 aggregate (orange) signals collected through imaging GV oocytes. Oocytes are circled using dashed white lines for clarity. Scale bar is 20 μm. *: p<0.05, **: p<0.01, ***: p<0.001. Error bars indicate standard error of mean. OCR: Oxygen consumption rate.

No differences were observed in ATP-production coupled respiration between the groups (Figure2E). Similar to basal respiration, proton leak was found to be significantly higher in CAPA-IVM GV-COCs compared to SO GV-COCs (Figure2F; p<0.001), while the latter group also exhibited significantly lower levels compared to baseline COCs (Figure2F; p<0.01). In vitro maturation also significantly decreased the proton leak in CAPA-IVM COCs (Figure2F; p< 0.001). As explained in Table 1, basal respiration is the net sum of processes capable of oxygen consuming that are used to produce ATP and proton leak [54]. Coupled respiration is the coupled part of respiratory oxygen flux that pumps the fraction of protons across the inner mitochondrial membrane which is utilized by the phosphorylation system to produce ATP from ADP and Pi. We detected higher coupling efficiency in SO GV-COCs compared to their in vitro counterparts (Figure2G; p<0.001), supporting the higher proportion of ATP production in this group compared to proton leak, despite comparable ATP-coupled respiration between the groups, indicating higher basal respiration is mainly due to increased proton leak. During in vivo growth with folligon, coupling efficiency also increased compared to baseline (Figure2G; p<0.01). Interestingly, coupling efficiency of in vitro matured COCs increased during IVM (Figure2G; p<0.001). As overall OCR are different between the groups, comparison between absolute spare respiratory capacity values would have been misleading, given it is a mathematical formula calculated by subtracting basal respiration from maximal respiration. Instead, we compared the spare respiratory capacity percentages, which showed a significant decrease in in vitro COCs compared to in vivo GV-COCs (Supplementary Figure 4A; p<0.05).

Non-mitochondrial oxygen consumption is presumed to increase when there is an increase of cytoplasmic ROS [55]. Non-mitochondrial oxygen consumption of CAPA-IVM GV-COCs was significantly higher compared to both baseline (p<0.05) and in vivo GV-COCs (p<0.001) (Figure2H). Similarly, CellROX staining of GV oocytes showed increased ROS content in oocytes following cytoplasmic maturation-both in vivo and in vitro-compared to baseline (Figure2J; p: baseline vs SO <0.001, baseline vs CAPA-IVM <0.0001). On the other hand, CAPA-IVM GV oocytes had significantly higher ROS levels compared to SO GVs, indicating the further ROS accumulating effect of the in vitro environment (Figure2J; p<0.05). While the mitochondrial membrane potentials of both SO and CAPA-IVM GV oocytes were significantly higher compared to baseline oocytes (both p values: <0.0001), no differences were detected between the two groups (Figure2K).

### Individual supplementation of lactate and super-GDF9 have limited effect on CAPA-IVM COC metabolism

We first compared the effect of lactate addition to pre-IVM only, and also to both steps of CAPA-IVM culture, to identify the best treatment protocol. Maturation rates and oocyte competency were comparable between the control and lactate supplemented group regardless of the CAPA-IVM step the COCs were cultured with lactate (Supplementary Figure 1C-E).

Next, we assessed the glycolytic activity of both CCs and oocytes with enzymatic assays following lactate supplementation. Specific enzymatic assays (PFK and LDH activities, pyruvate and lactate concentrations) were selected based on the upregulated glycolysis observed in the previous study [37]. Lactate supplementation during the pre-IVM step of CAPA-IVM culture did not cause any changes in CCs for any of the studied enzymes or metabolites, even though some values were almost halved during the culture (Figure3 A-D). Within MII-CCs, pyruvate and lactate levels, as well as PFK and LDH activities between the conditions were comparable to the GV stage (Figure3 A-D). Similarly, media lactate supplementation during pre-IVM or both steps did not affect CC pyruvate levels or PFK activity since they were comparable between GVs and both lactate MIIs. LDH activity (Figure3C; p: 0.0051) and lactate concentration (Figure3D; p: 0.0098) of MII-CCs increased significantly compared to GV-CCs when lactate was added throughout the CAPA-IVM culture period, instead of only in the pre-IVM step.

Pyruvate levels of CCs were significantly higher compared to oocytes from control GV- and MII-COC, as well as lactate supplemented GV-COCs (Supplementary Figure 2A). On the other hand, PFK activity of CCs compared to oocytes were significantly higher in all of the analyzed groups (Supplementary Figure 2B). LDH activity of CCs and oocytes were comparable only in lactate supplemented GV-CCs (Supplementary Figure2C). Significant differences between MII stage CC and oocyte lactate concentrations were observed when culture media was supplemented with lactate either during pre-IVM or both steps of CAPA-IVM (Supplementary Figure 2D).

GDF9 supplementation of pre-IVM media did not affect pyruvate and lactate concentrations of GV-CCs (Figure 3E and H). On the other hand, GDF9 supplementation significantly increased LDH activity in GV-CCs compared to the control group (Figure 3G; p <0.0001). Pyruvate concentrations and PFK activities of MII-CCs were comparable regardless of the media supplementation (Figure 3E and F). During GV-to-MII transition a significant decrease in pyruvate was detected, both in the control group and GDF9 supplemented group MII-CCs compared to GV-CCs (Figure 3E; p values: 0.0038 and 0.0003, respectively). On the contrary, LDH activity was significantly higher in the control group MII-CCs compared to GV-CCs (Figure 3G; p: 0.0327).

**Figure 3:**
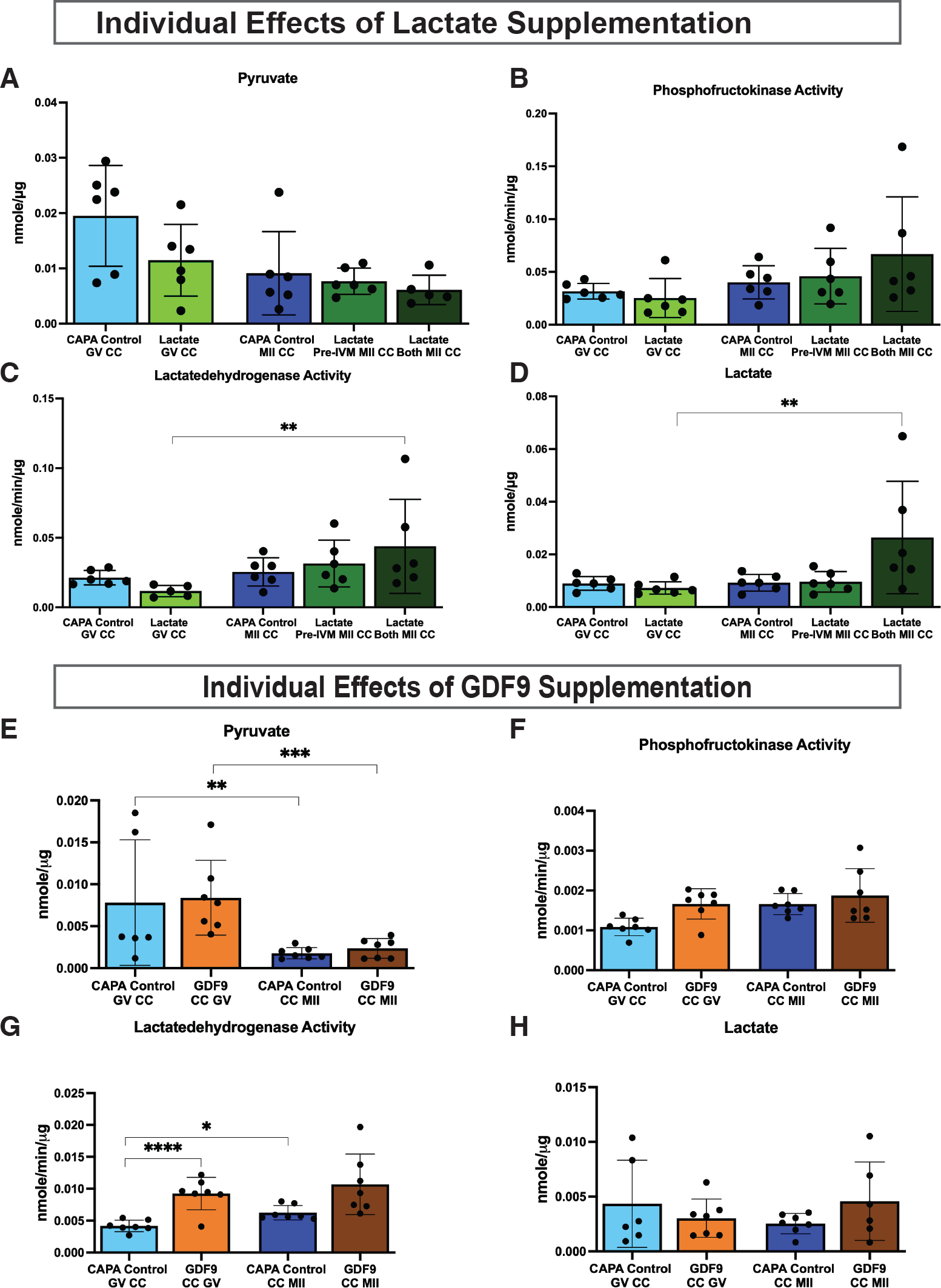
Individual effects of lactate **(A,B,C,D)** and super-GDF9 **(E,F,G,H)** supplementation during CAPA-IVM culture were assessed through enzymatic assays in cumulus cells. Each dot represents one biological replicate. *: p<0.05, **: p<0.01, ***: p<0.001, ****: p<0.0001. Error bars indicate standard deviation.

Pyruvate levels of CCs from both GV-COC conditions were significantly higher than in oocytes (Supplementary Figure 3A). Both PFK and LDH activities of CCs compared to oocytes were significantly higher in all of the analyzed groups (Supplementary Figure 3B and C). CC lactate concentrations were higher than those in oocytes of GV-COCs from the control group, and GDF9 supplemented MII-COCs (Supplementary Figure 3D).

### Combined lactate+super-GDF9 supplementation does not restore the boost in pre-IVM respiration

Oocyte maturation rates and IVF results, reflecting oocyte competence, of the groups are provided in Table 2. No differences were detected regarding the mature oocyte rate, D2 cleavage rate or D5 blastocyst rate, between any of the groups. When measuring the diameters of MII oocytes, we found that oocytes from the lactate only supplemented (mean diameter: 71.7±1.6 μm, n: 68) and GDF9 only supplemented (mean diameter: 71.6±1.9 μm, n: 52) groups were larger in size compared to the control group (mean diameter: 70.7±2.1 μm, n: 66; respective p values are 0.0064 and 0.0357), while they were comparable to the lactate+GDF9 combined group.

After cytoplasmic maturation (pre-IVM), both maximal and basal respirations significantly increased in groups receiving GDF9 (both p<0.01) and lactate+GDF9 supplementation (basal: p<0.001; maximal: p<0.01) compared to control GV-COCs (Figure 4C and D). Combining lactate supplementation with GDF9 significantly increased the basal respiration in this group compared to lactate only supplementation (Figure 4C, p<0.001). Increase in the ATP coupled respiration in GDF9+lactate supplemented group led to a significant difference in this function compared to control group (Figure 4F, p<0.01). Non-mitochondrial oxygen consumption was significantly higher in GDF9+lactate supplemented group compared to all other in vitro grown GV-COCs (Figure 4H, all p<0.001). Control and lactate only supplemented groups had significantly lower proton leaks compared to both GDF9 only (vs: control p<0.01, lactate p<0.05) and GDF9+lactate (vs: control p<0.001, lactate p<0.05) supplemented groups (Figure 4G).

**Figure 4:**
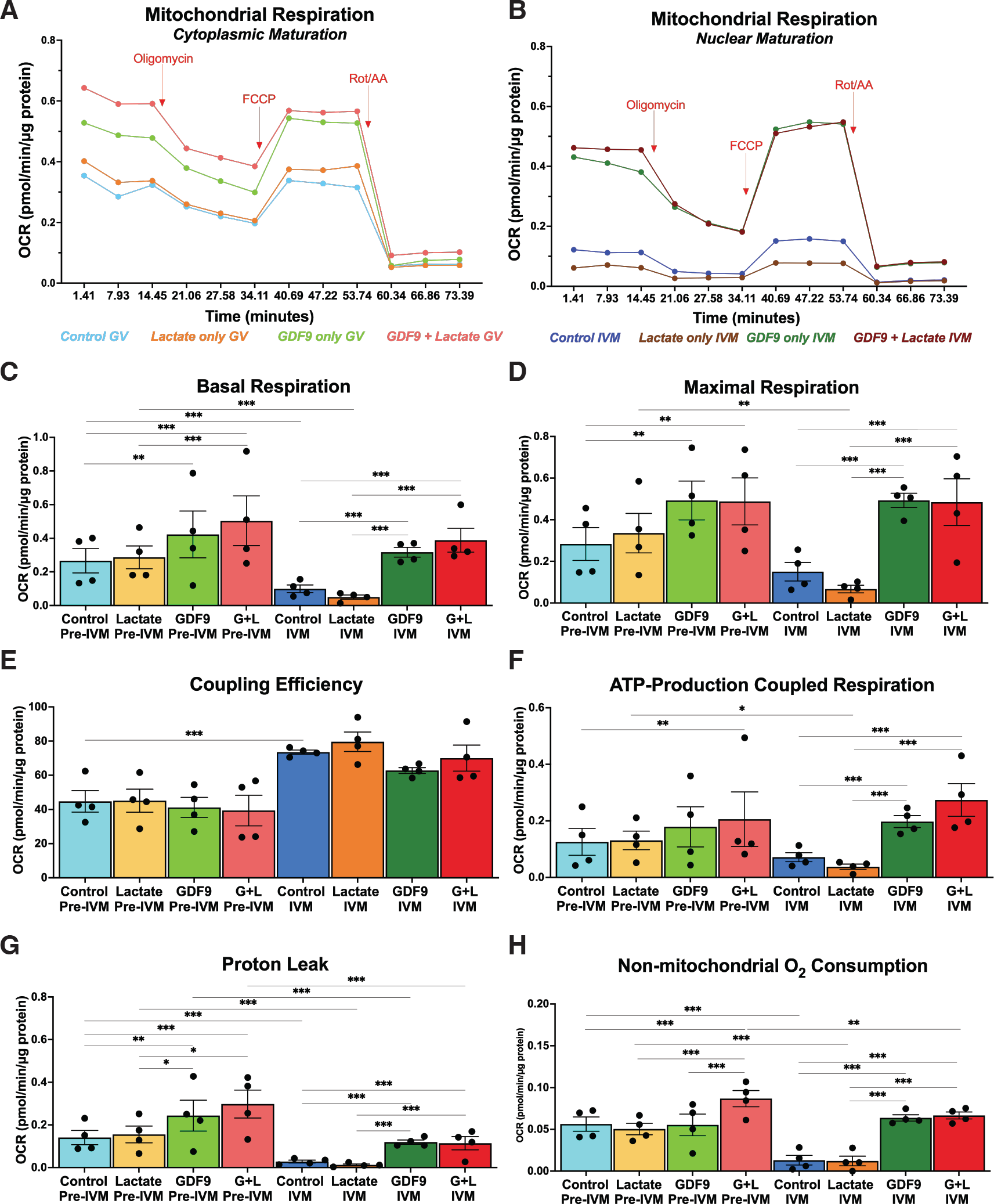
Mitochondrial function was compared within CAPA-IVM cultured GV- and MII-COCs. through real-time metabolic analysis with Mito Stress Test (Seahorse Analyzer). *: p<0.05, **: p<0.01, ***: p<0.001. Error bars indicate standard error of mean. L: Lactate. G: Super-GDF9. OCR: Oxygen consumption rate.

The pattern observed in maximal and basal respiration rates in COCs after pre-IVM was also detected after IVM, as both functions were significantly higher after IVM in GDF9 only and lactate+GDF9 supplemented groups compared to control (Figure 4C and D, all p values <0.001). Likewise, after IVM, maximal and basal respiration of lactate supplemented group were significantly lower compared to GDF9 only and lactate+GDF9 supplemented groups (Figure 4C and D, all p values <0.001). Furthermore, within MII-COCs GDF9 only and lactate+GDF9 supplemented groups had significantly higher ATP-production coupled respiration, proton leak and non-mitochondrial oxygen consumption compared to both control and lactate supplemented groups (Figures 4F-H, all p values <0.001). These aforementioned functions were comparable between control vs. lactate only and GDF9 only vs. lactate+GDF9 supplemented group MII-COCs. Coupling efficiency (Figure 4E) and spare respiratory capacity (Supplementary Figure 4B) within MII-COCs were comparable.

When comparing changes during the pre-IVM to IVM transition, both basal and maximal respiration were decreased in the COCs of lactate only supplemented group (p<0.001 and p<0.01, respectively), while this decrease was limited to basal respiration in the control group (p<0.001) (Figure 4C and D). Following nuclear maturation, the coupling efficiency was significantly higher in the control group (p<0.001, Figure 4E), and a significant decrease in ATP-production coupled respiration was observed in lactate only supplemented group (p<0.05, Figure 4F). A global decrease was observed in proton leak following nuclear maturation in all MII-COCs (all p<0.001, Figure 4G). Similarly, non-mitochondrial oxygen consumption significantly decreased during the GV-to-MII transition in the control (p<0.001), lactate only (p<0.001) and lactate+GDF9 supplemented groups (p<0.01) (Figure 4H). No changes in the spare respiratory capacity were observed (Supplementary Figure 4B).

Finally, no differences were detected in ROS accumulation and mitochondrial membrane potential between CAPA-IVM oocytes, whether the culture media was supplemented with lactate and/or GDF9 or not (Figure 5).

**Figure 5:**
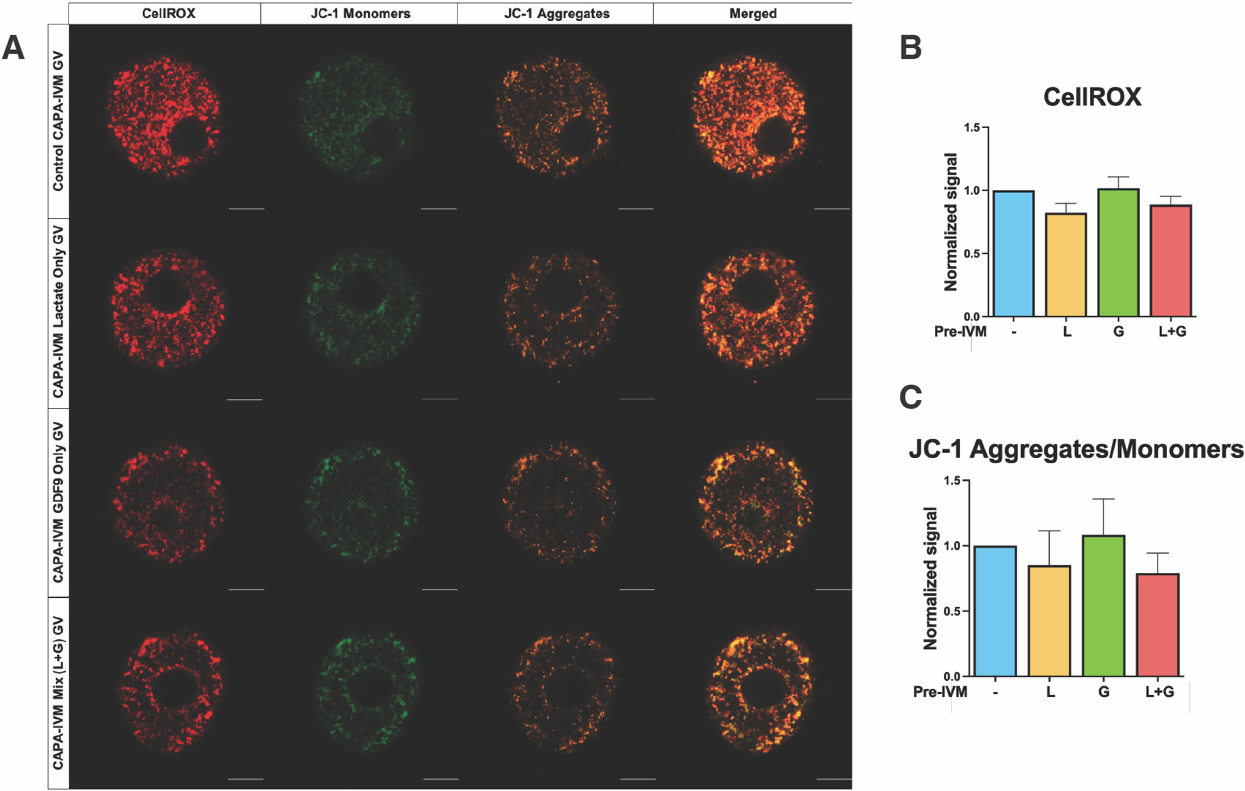
ROS accumulation and mitochondrial membrane potential were compared within CAPA-IVM cultured GV oocytes. **(A)** Panel illustrates CellROX (red), JC-1 monomer (green) and JC-1 aggregate (orange) signals collected through imaging GV oocytes. Scale bar is 20 μm. **(B)** shows average CellROX signal and **(C)** shows mitochondrial membrane potential, both data collected from three separate experiments. L: Lactate. G: Super-GDF9.

### Low oxygen tension during pre-IVM restores basal respiration profile of CAPA-IVM COCs

Oocyte maturation rates and IVF results, indicating oocyte competence, of the groups are provided in Table 2. No differences were detected regarding mature oocytes rate, MII oocyte diameter, D2 cleavage rate or D5 blastocyst rate between groups (Table 2).

**Table 2:** Effect of lactate and/or super-GDF9 addition (Experiment II) and pre-IVM oxygen tension (Experiment III) on CAPA-IVM oocyte maturation and competence. MII rate was assessed through denuding 72-92 expanded COCs. MII oocyte size was determined through measuring 56-73 MII oocytes. COCs for both assessments were obtained from in total 76 mice (three individual experiments). IVFs were performed via inseminating 64-98 expanded COCs (three individual experiments) obtained from 60 mice. Results represented as mean ± SD.

| Experiment II |  |  |  |  |
| --- | --- | --- | --- | --- |
|  | MII oocyte (%) | MII oocyte size (μm) | Cleavage (%) | Blastocyst/Cleaved % |
| Control | 77.7 ± 3.5 | 70.7 ± 2.1<sup>a</sup> | 53 ± 19.1 | 78 ± 5 |
| Lactate only | 80.3 ± 7.1 | 71.7 ± 1.6<sup>b</sup> | 49.7 ± 16.9 | 80.7 ± 18.1 |
| GDF9 only | 73 ± 6 | 71.6 ± 1.9<sup>bc</sup> | 59.3 ± 9.5 | 62 ± 27.8 |
| Lactate + GDF9 | 75.7 ± 8.1 | 71.3 ± 1.6<sup>abc</sup> | 55.3 ± 30 | 70.3 ± 36.1 |
| Experiment III |  |  |  |  |
|  | MII oocyte (%) | MII oocyte size (μm) | Cleavage (%) | Blastocyst/Cleaved % |
| Control Atm O<sub>2</sub> | 77.7 ± 3.5 | 70.7 ± 2.1 | 57 ± 16.4 | 79.7 ± 2.9 |
| Control Low O<sub>2</sub> | 78.3 ± 6.8 | 70.9 ± 1.9 | 38 ± 19.9 | 73.3 ± 30 |

Mitochondrial respiration profiles showed that when COCs were cultured in low oxygen during pre-IVM, their profile resembled SO GV-COCs more than CAPA-IVM control GV-COCs cultured in atmospheric oxygen (Figure 6A). On the other hand, mitochondrial respiration of the same group of COCs were lower compared to both SO and CAPA-IVM control groups at MII stage (Figure 6B). While compared to SO GV-COCs, CAPA-IVM control GV-COCs cultured in atmospheric oxygen exhibited significantly higher basal respiration (Figure 6C, p<0.001), maximal respiration (Figure 6D, p<0.01), proton leak (Figure 6E, p<0.001) and non-mitochondrial oxygen consumption (Figure 6F, p<0.001), the levels of these functions were comparable between SO-GV-COCs and CAPA-IVM control GV-COCs cultured in low (5%) oxygen (Figures 6C-F). Within CAPA-IVM conditions no differences were detected besides from significantly lower non-mitochondrial oxygen consumption in GV-COCs cultured in low oxygen (Figure 6G, p<0.001). Coupling efficiency (both p<0.001) and spare respiratory capacity % (both p<0.05) were both significantly higher in SO GV-COCs compared to both in vitro conditions (Figure 6H, Supplementary Figure 4C).

**Figure 6:**
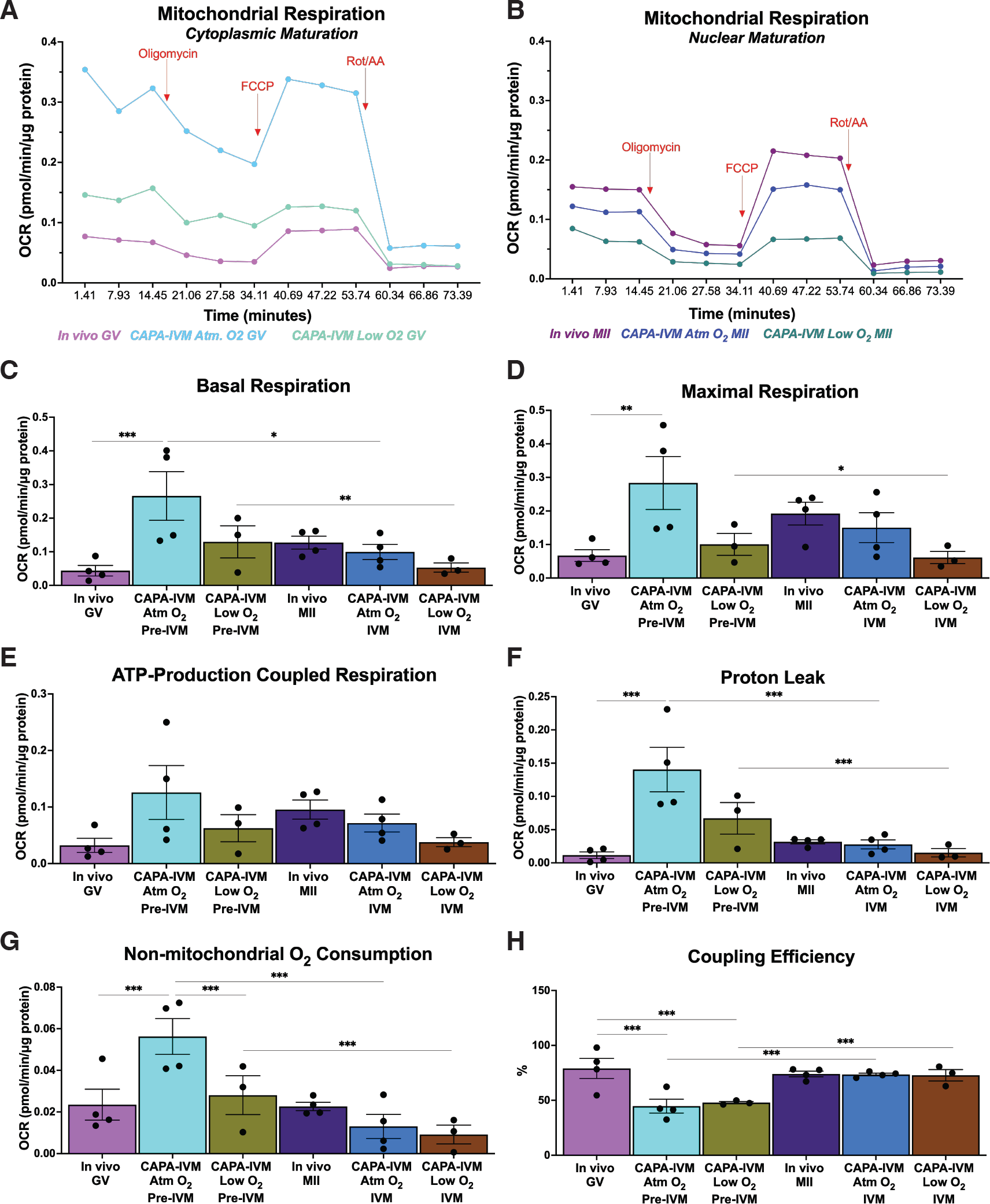
Mitochondrial function was compared between SO COCs, and CAPA-IVM COCs cultured under atmospheric oxygen vs low oxygen during pre-IVM, through real-time metabolic analysis with Mito Stress Test (Seahorse Analyzer). *: p<0.05, **: p<0.01, ***:p<0.001. Error bars indicate standard error of mean. Atm: Atmospheric. OCR: Oxygen consumption rate.

There were no differences within the MII-COCs of the three conditions. In vitro nuclear maturation decreased basal respiration (Figure 6C, p_20%Oxygen_<0.05, p_5%Oxygen_<0.01), proton leak (Figure 6F, both p<0.001), non-mitochondrial oxygen consumption (Figure 6G, p<0.001) and increased the coupling efficiency (Figure 6H, p<0.001) in both in vitro conditions, regardless of the oxygen tension they were exposed during pre-IVM. The decrease in the MII-COCs maximal respiration was significant in the CAPA-IVM group that was cultured under 5% oxygen tension during pre-IVM (figure 6D, p<0.05).

No differences were detected in ROS accumulation and mitochondrial membrane potential between CAPA-IVM oocytes whether they were cultured in atmospheric oxygen or low oxygen during the pre-IVM culture period (Supplementary Figure 5).

## DISCUSSION

Through recent advancements partially revealing the complexity of oocyte maturation, and concomitantly, introduction of bi-phasic IVM systems, IVM oocytes’ developmental competence has been successfully enhanced. CAPA-IVM is one such bi-phasic IVM protocol, has a proven superiority over standard IVM protocols in humans, with improved maturation and pregnancy rates [10,12–14]. Despite this success, the high attrition rates from mature oocytes to good quality embryos is still keeping the efficiency of CAPA-IVM at lower levels compared to conventional IVF. Recently we have shown perturbances in CC glucose metabolism of CAPA-IVM COCs that are inflicted during pre-IVM culture, leading to precociously increased glycolytic activity [37]. Considering the metabolic needs and the microenvironment of the maturing small-/mid-antral COCs, we hypothesized that further optimization of CAPA-IVM could be possible through focusing on adjusting culture media nutrient concentrations and/or oxygen tension [37].

Analyzing real-time OCR, we found that following Folligon injection, mitochondrial respiration as well as the basal and maximal OCR of the SO GV-COCs were decreased compared to baseline COCs, which was not observed in in vitro COCs. This dramatic difference between SO GV-COCs and CAPA-IVM GV-COCs was observed also in the basal and maximal respiration rates, showing that OCR was significantly higher in in vitro cultured GV-COCs compared to SO. Considering the essential bi-directional communication between CCs and oocytes [33,41], and taken together with our previous results indicating a boost in CC glycolysis during pre-IVM [37], current results show that it is the increased OXPHOS that stimulates precocious CC glycolysis, since pyruvate oxidation by oocytes cannot be completed without substrates provided by CCs [31,56].

Basal respiration is the net sum of processes capable of consuming oxygen which are ATP-production coupled respiration and proton leak [54]. We found that in CAPA-IVM GV-COCs, proton leak is significantly higher compared to SO GV-COCs, while no difference was observed for the ATP-production coupled respiration. Proton leak reflects the level of protons migrating to the mitochondrial matrix without resulting in ATP production [57]. Furthermore, CellROX staining of GV oocytes showed that in vivo cytoplasmic maturation increases the abundance of ROS even though proton leak decreases alongside with increased coupling efficiency. Although ROS are in general perceived as damaging by-products of OXPHOS, they are also important regulatory signaling molecules responsible for growth and development [58], a finding supported through our current results as well. However, given the significantly higher CellROX signal in the CAPA-IVM GV oocytes compared to SO oocytes, it is plausible to say that the pre-IVM culture is inflicting oxidative stress.

We did not observe any significant differences within respiration profiles of MII-COCs, indicating that despite the perturbances were inflicted during pre-IVM culture, the bioenergetic profile of COCs was able to resume to normo-patterns during IVM. Even though, in vtiro GV-to-MII transition was marked by a decrease in mitochondrial, basal and maximal respiration, supporting the findings by Scantland et al. [59], the coupling efficiency and the percentage of basal respiration dedicated for ATP-production increased (not shown), probably in an attempt by the oocyte to prevent ATP deficit. During GV-to-MII transition CAPA-IVM COCs non-mitochondrial oxygen consumption significantly decreased. This function is presumed to increase when there is an increase of cytoplasmic ROS [55]. Similarly, its decrease can possibly be explained as removal of the ROS from the environment. As proton leak also decreased and became comparable between MII-COCs of in vitro and in vivo, this detected decrease in non-mitochondrial oxygen consumption could be another indication for restoration of normal bioenergetic patterns during the IVM phase.

Lactate supplementation to culture media did not induce any significant changes in CC glycolysis. When media was supplemented with lactate during both steps of CAPA-IVM, lactate concentrations and LDH activity increased in CCs, without changing pyruvate concentration or PFK activity. While this is indicative of lactate being taken up from the environment by the CCs and metabolized, comparable mitochondrial respiration in COCs supports the findings of Dumollard et al. indicating that lactate-derived pyruvate cannot be used by oocytes to produce ATP [30]. GDF9 only supplementation to culture media slightly increased the activity of glycolytic enzyme PFK in CCs, although not significantly. On the other hand, while GDF9 supplementation led to a significant increase in LDH activity, we did not observe any difference in CC pyruvate and lactate levels. Given that mitochondrial respiration increased in GDF9 supplemented CAPA-IVM COCs, even though we did not observe a significant increase in CC PFK activity, it is clear that more substrates were delivered to the oocyte for ATP production by the CC compartment, as oocyte and CC metabolisms are interdependent [41,60,61].

Real-time metabolic analysis of COCs showed that none of the supplement combinations were able to restore the in vivo COC’s mitochondrial respiration profile. On the other hand, COCs of lactate only supplemented group exhibited the only significant decrease in the ATP-production coupled respiration during GV-to-MII transition. Even though the ratio of ATP-coupled respiration to proton leak was comparable within all CAPA-IVM MII-COCs, the former decrease might result in decreased ATP concentrations within oocytes. Furthermore, results indicated that GDF9 is the principal substance inflicting changes in oocyte’s oxygen consumption, since GDF9 only and lactate+GDF9 mix supplemented COCs exhibited comparable OCR for all analyzed components, both at GV and MII stages. The only difference between GDF9 only and lactate+GDF9 mix supplemented COCs was observed in non-mitochondrial oxygen consumption at the end of pre-IVM, which was significantly higher in the group supplemented with the mix compared to all other in vitro groups. Yet, there was not a significant difference in ROS production and mitochondrial function between GV-COCs, nor in oocyte competency following IVF after CAPA-IVM. In leukocytes, the increase in non-mitochondrial OCR is attributed to the increase of inflammatory enzymes [62]. COC expansion and ovulation during oocyte maturation have been compared to an immune-like response, with similar characteristics to inflammation [63]. IVM CCs generally have aberrant expression of genes from the aforementioned processes [64], but OSF addition during CAPA-IVM culture significantly regulates expression of genes involved in CC expansion to a level comparable to in vivo matured COCs [42]. While we did not observe any difference in the mucification between GDF9 only vs lactate+GDF9 supplemented groups, it is clear that using these supplements together is augmenting their effects on non-mitochondrial OCR in GV COCs. Regardless, this difference in non-mitochondrial OCR calls for further studies for clarification of the observed effect. [68][69] Lactate is the end-product of lactic acid fermentation, with a known role of inhibiting the activity of the glycolysis enzyme PFK in several tissues [38]. Yet, it failed to act as a glycolysis-limiting nutrient as expected during the pre-IVM step. Consequently, we concluded that regardless of the culture media supplementation, it is the pre-IVM culture environment itself that is boosting metabolic activity in COCs. The direct relationship between culture environment oxygen concentration and the ATP content of IVM oocytes has been shown by Hashimoto et al. [46]. Oxygen concentration of the reproductive tract varies between 2%-9% [44,45] which is significantly lower than atmospheric oxygen concentrations. However, the optimal oxygen microenvironment to use during IVM is still one of the most debated topics of optimization strategies, as studies have reported contradictive results. Banwell et al. indicated that changing oxygen concentrations during mouse IVM does not affect oocyte maturation, fertilization, or embryo development rates [65]. On the other hand, Preis et al. showed improved oocyte competence following IVM at 5% oxygen [66]. In bovine IVM, 5% oxygen tension reduced the number of mature oocytes, but those that matured exhibited better embryonic developmental capacity compared to the oocytes matured under 20% oxygen [46]. In the current study, given no differences were observed in COC metabolic activity following media nutrient supplementation, we cultured the COCs in a low oxygen environment during pre-IVM to physically limit their access to oxygen and tested oocyte competency and COC metabolism.

COC metabolic profile changed strikingly upon culturing them at low oxygen tension during pre-IVM. We found that GV-COCs cultured under 5% oxygen during pre-IVM had decreased mitochondrial respiration. Both basal and maximal respiration levels, as well as the proton leak of the 5% oxygen group were comparable to both SO-GV and in vitro COCs cultured under 20% O_2_. On the other hand, culturing the COCs under 5% oxygen restored the non-mitochondrial OCR in this group. Surprisingly, we did not detect any difference in GV oocyte ROS levels and mitochondrial membrane potential after reducing the culture oxygen tension. In embryos, higher metabolic activity is related to lower developmental potential due to molecular and cellular damage inflicted by ROS accumulation [67–69]. Elevated oxygen concentrations increase ROS generation [70], which leads to severe damage in DNA. In fact, the highest DNA damage rates are found in the most metabolically active embryos [71]. Upon sensing external stimuli, cells normally respond to maintain homeostasis through regulating the expression of several genes [72]. However, transcriptional machinery is regulated differentially in oocytes compared to somatic cells, as mRNA transcription ceases by the time the oocyte is ready for nuclear maturation [73,74]. Following the resumption of meiosis, maternally stored mRNAs are translated to support the zygote until activation of their genome [75]. Therefore, in addition to higher ROS accumulation due to hyper-active metabolism, oocytes might not be equipped to maintain homeostasis through stimulating transcriptomic changes upon encountering hyperoxia during CAPA-IVM, ultimately leading to lower embryo quality.

Both GV and MII oocytes exhibit an atypical hooded mitochondrial ultrastructure, with fewer cristae, with a reduced capacity for OXPHOS, compared to other cell types [59,76–78]. In contrast, the energy demands of an oocyte during cytoplasmic maturation is massive given the essential biological transition including cytoskeleton remodeling, organelle reorganization and nuclear reprograming [79]. Essentially, a cell’s energy demand is the driving force for ATP production, which is challenged under stress conditions. The fact that we observed significant differences in spare respiratory capacity % in GV-COCs from both in vitro conditions compared to SO GV-COCs indicates that mitochondrial flexibility is not preserved during stress-inducing pre-IVM culture. On the other hand, the spare respiratory capacity ratio was comparable between the MII-COC. Moreover, considering the assessed endpoints for oocyte competence between COCs cultured under 5% vs 20% O_2_ during pre-IVM did not reflect any changes, the current slightly lower mitochondrial respiration profile of the MII-COCs cultured under 5% oxygen during pre-IVM is not alarming. Recently, the adenosine salvage pathway has been suggested as an alternative and complementary ATP production pathway to prevent energy deficit during oocyte maturation [59,80]. This two-step reaction that produces ATP through accumulated cAMP could be the reason for comparable oocyte competence between the two in vitro groups. However, given that mitochondria are inherited maternally, oocytes with dysfunctional mitochondria might still lead to deficiencies post maturation, and cause lower rates of good quality embryos in IVM systems. Thus, it is necessary to focus on improving spare respiratory capacity of GV-COCs cultured in vitro at 5% oxygen during pre-IVM and combine it with improving basal respiration rates and balancing ROS levels of MII-COCs during the IVM step.

Our study did not highlight any improvement in oocyte competence, with the current assessment criteria, following media supplementation of lactate and/or GDF9. Our current unstimulated pre-pubertal mouse model is successful in imitating the low competence human oocytes obtained from 2-8 mm follicles after minimal FSH stimulation. Our long-term aim is to assess the success of the advancements in our mouse CAPA-IVM protocol by embryo implantation potential, live birth rates and newborn health. On the other hand, one limitation of the model is the absence of any infertility etiology that interferes with the detection of short-term improvements in oocyte competence [42]. Testing our current findings in a PCOS mouse model would be very interesting, given patients with PCOS have a distorted metabolic profile compared to normo-ovulatory patients which causes increased oxidative stress in oocytes [81]. A ROS regulatory role of CAPA-IVM has already been proposed through observed changes in CC gene expression [13]. In this study, even though reduced oxygen tension was successful in improving the bioenergetic profile, it did not change the intensity of accumulated ROS in GV oocytes. Hence, antioxidant supplementation could also be utilized to improve ROS accumulation during CAPA-IVM under low oxygen concentrations.

To the best of our knowledge, this is the first study reporting the respiratory profiles of mouse COCs with real-time metabolic analysis, during bi-phasic IVM and during in vivo maturation. Results show that pre-IVM culture is boosting OXPHOS in GV-COCs and inflicting oxidative stress in GV oocytes. Neither individual nor combined supplementations of lactate and GDF9 are effective in limiting pre-IVM CC glycolysis. However, the reduction of oxygen tension during pre-IVM, is a promising culture strategy to regulate OCR and metabolism of CAPA-IVM COCs, paving the way for future investigations in the human CAPA-IVM system.

## ACKNOWLEDGEMENTS

The authors thank Aimilia Zisiadi for her contribution in preparation of in vivo samples.

