## Supplemental figures 1-6 for "Effects of lactate, super-GDF9 and low oxygen tension during biphasic in vitro maturation on the bioenergetic profiles of mouse cumulus-oocyte-complex"

### SUPPLEMENTARY FIGURES

#### Supplementary Figure 1

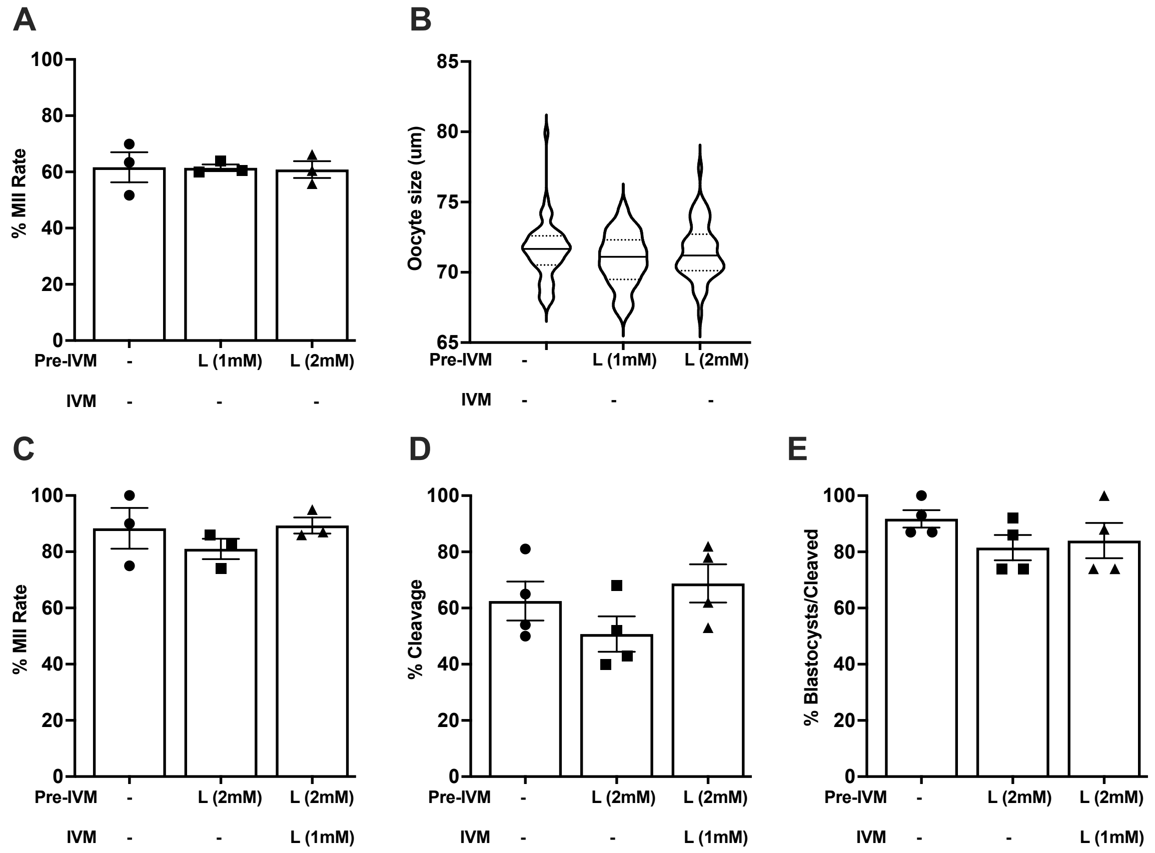


**Supplementary Figure 1: (A and B)** are showing the results of the dose finding study for lactate. Neither of the doses indicated any disruption. Considering the reports from literature, we decided to use 2 mM in our experiments. **(C-E)** are showing the mature oocyte rates and IVF results following the initial test of lactate supplementation to CAPA-IVM media. Given no significant differences were seen between experimental groups, we decided to supplement lactate both steps of CAPA-IVM as it is also present in the follicular fluid and reproductive tract throughout the menstrual cycle.

### Supplementary Figure 2

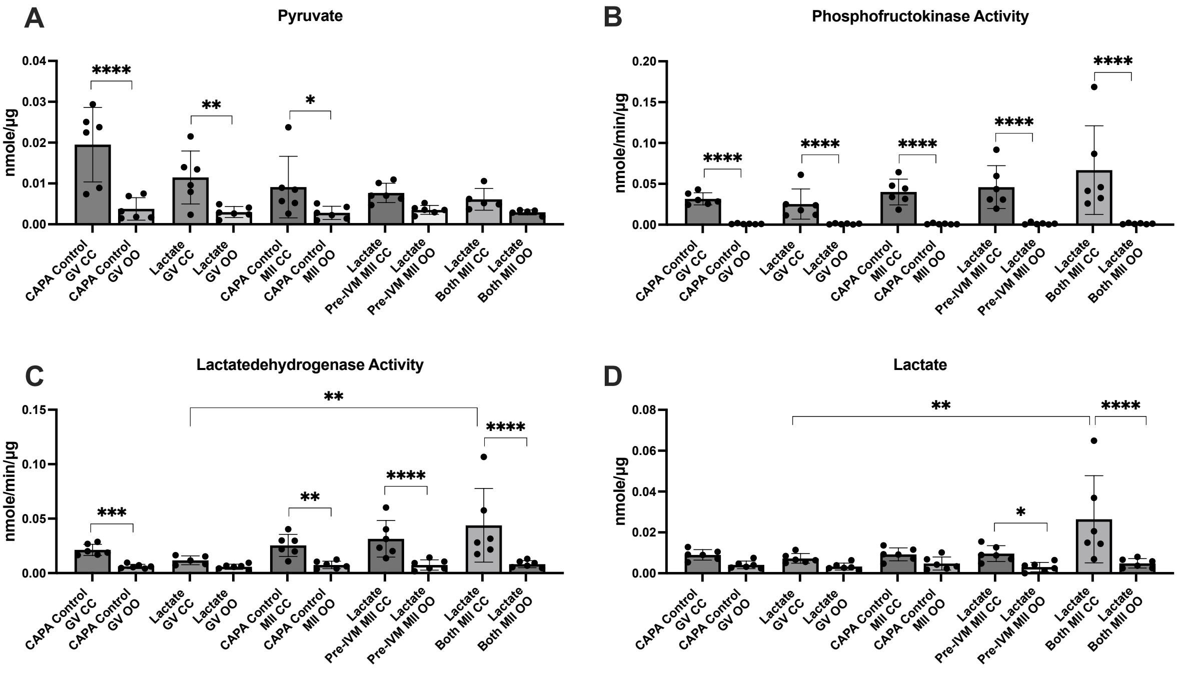


**Supplementary Figure 2:** Graphs showing individual effect of lactate supplementation both with CCs and oocytes. *: $p<0.05$, **: $p<0.01$, ***: $p<0.001$, ****: $p<0.0001$. Error bars indicate standard deviation.

### Supplementary Figure 3

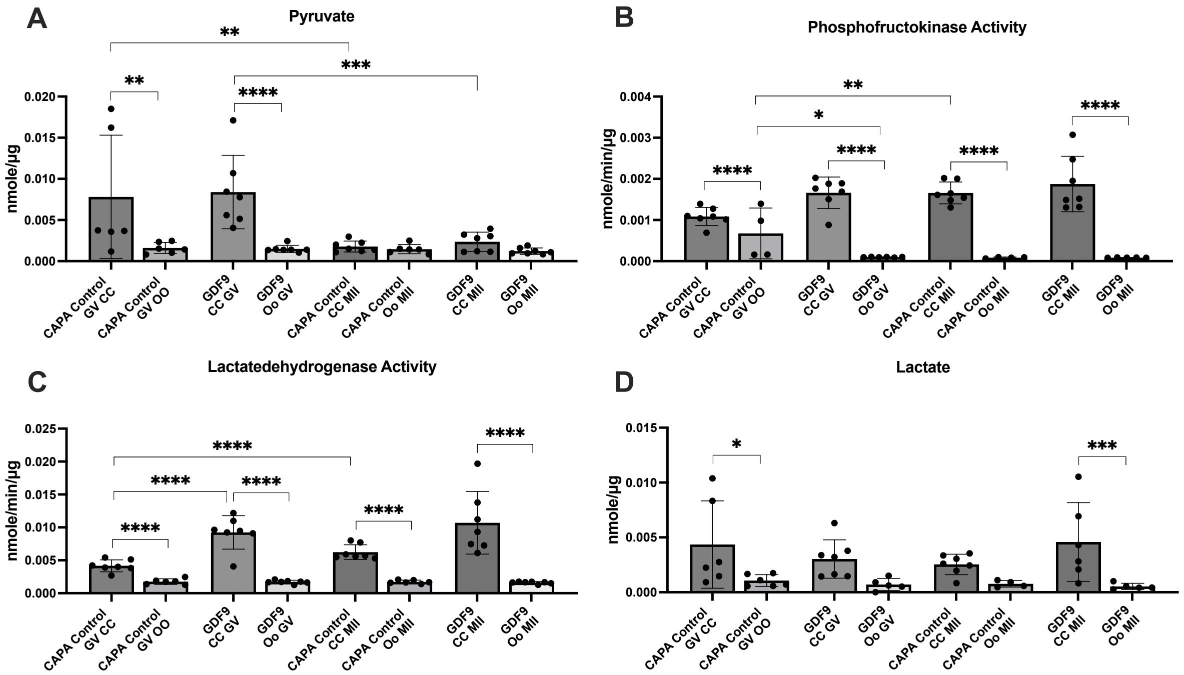


**Supplementary Figure 3:** Graphs showing individual effect of super-GDF9 supplementation both with CCs and oocytes. *: p<0.05, **: p<0.01, ***: p<0.001, ****: p<0.0001. Error bars indicate standard deviation.

### Supplementary Figure 4

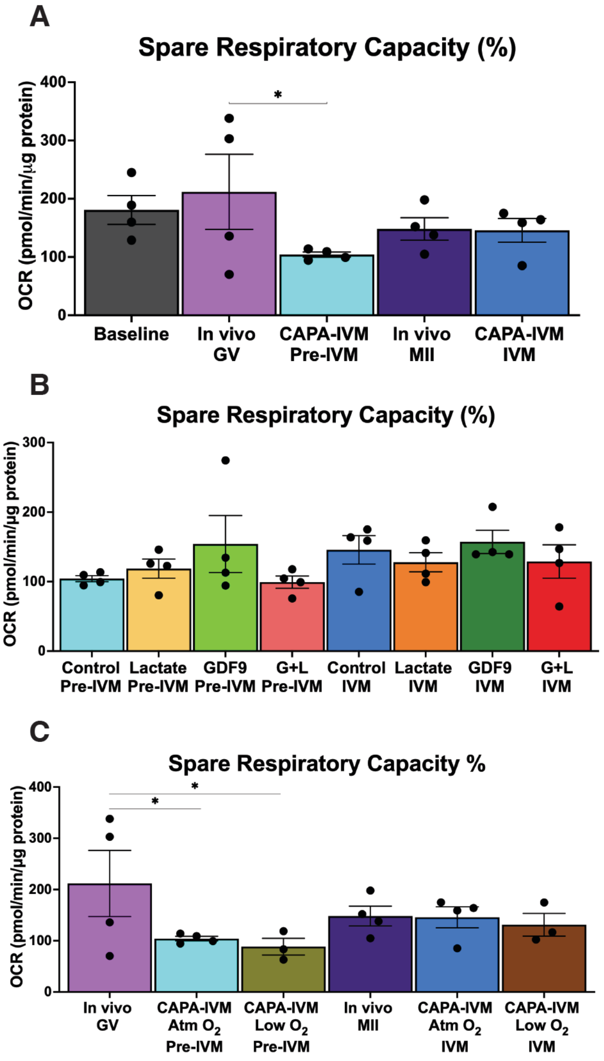


**Supplementary Figure 4:** Spare respiratory capacity % of (A) in vivo vs CAPA-IVM matured COCs, (B) CAPA-IVM cultured GV- and MII-COCs with different supplementations, (C) in vivo COCs, and CAPA-IVM COCs cultured under atmospheric oxygen vs low oxygen during pre-IVM. *: $p<0.05$. Error bars indicate standard error of mean L: Lactate. G: Super-GDF9. Atm: Atmospheric. OCR: Oxygen consumption rate.

**Supplementary Figure 5**

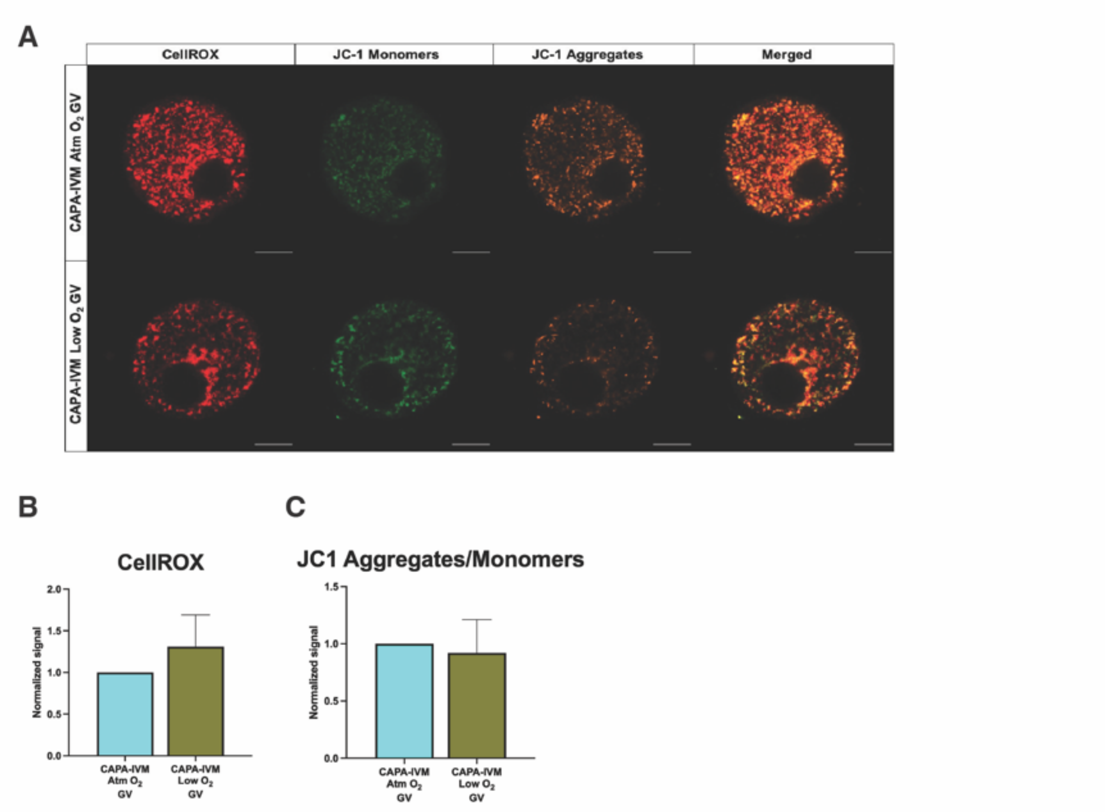


**Supplementary Figure 5:** ROS accumulation and mitochondrial membrane potential were compared within CAPA-IVM cultured GV oocytes under atmospheric (Atm) oxygen and 5% (low) oxygen. **(A)** Panel illustrates CellROX (red), JC-1 monomer (green) and JC-1 aggregate (orange) signals collected through imaging GV oocytes. Scale bar is 20 μm. **(B)** shows average CellROX signal and **(C)** shows mitochondrial membrane potential, both data collected from three separate experiments.

### Supplementary Figure 6

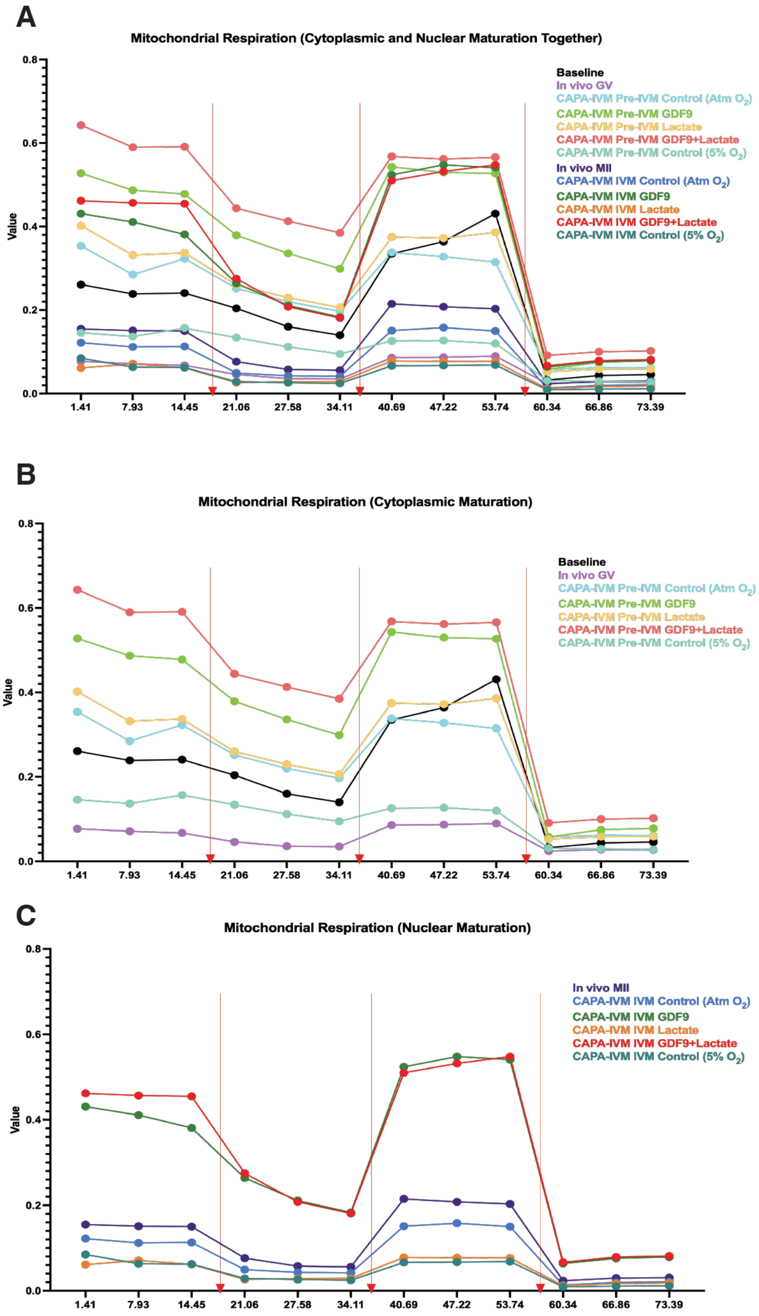


**Supplementary Figure 6:** Mitochondrial respiration profiles of each group are represented together (A) regardless of maturation and/or culture step, (B) after pre-IVM and (C) after IVM.
